# Phylogenetically and ecologically delimited species pool affect the inference of age-dependent extinction

**DOI:** 10.1101/2024.10.11.616654

**Authors:** Salatiel Gonçalves-Neto, Tiago B. Quental

## Abstract

According to Van Valen’s seminal work (1973), extinction occurs at a constantly stochastic rate within ecologically homogeneous groups or adaptive zones, giving long and short-lived species equal chances of extinction. Van Valen highlighted the difficulty in defining and identifying the species pool within an adaptive zone, but fundamentally viewed it through ecological factors. Most studies have used taxonomically or phylogenetically defined species pools to test the “Law of Constant Extinction.” Here, we investigate how different species pools defined by phylogeny or ecology influence the inference of age-independent extinction. Using the Canidae fossil record and a Bayesian framework, we show that species pools defined by phylogeny or ecology exhibit different age-dependent extinction dynamics. The age-dependent extinction (ADE) signal varies depending on the species pool choice, time window, and taxonomic level. Within phylogenetic species pools, we observe mixed evidence for ADE, with both positive—older species more likely to go extinct (Hesperocyoninae and Borophaginae)—and negative—younger species more likely to go extinct (Caninae)—trends. Combining subfamilies into a family-level analysis yields weak evidence for ADE or strong support for age-independent extinction, depending on the period analyzed. Within ecologically defined species pools, hypercarnivores show strong evidence for positive ADE, whereas non-hypercarnivores show signals akin to age-independent extinction. Phylogenetic pools with more hypercarnivores tended to show evidence of positive ADE, while those with fewer tended toward negative ADE. These findings emphasize that the choice of species pool significantly influences observed age-dependent extinction dynamics and that ecology impacts the regime of age-dependent extinction.

## Introduction

Based on linear survivorship curves (presented on a semi-logarithmic scale), Van Valen (1973) asserted that extinction occurs at a consistently stochastic rate within an adaptive zone or an ecologically homogeneous taxonomic group, resulting in an equal chance of species going extinct at any given point in time. This pattern led him to suggest that the probability of extinction was unrelated to the age of a lineage, a phenomenon that seems to be nearly ubiquitous across all forms of biodiversity when looking at individual clades and was termed “The Law of Constant Extinction” (Van Valen, 1973). He introduced the Red Queen hypothesis as a mechanism to account for his “law” (Van Valen, 1973), relying on the concept of adaptive zones coined by G.G. Simpson some years earlier (Simpson, 1944).

An adaptive zone could be characterized as: “*the sum of the physical and biotic situations that an organism or a group of ecologically and/or evolutionarily related entities lives and evolves in. This can be thought of as the niche of a higher taxon. It is itself evolvable and particular adaptive zones might be unfilled by organisms at a given time*” (Liow *et al*., 2011). Hence, in accordance with Van Valen’s Red Queen hypothesis, within a given adaptive zone, lineages must perpetually adapt to match the ever-changing environment. It’s within this adaptive zone that the “Law of Constant Extinction” becomes apparent. At the core of Van Valen’s argument is the notion that the total amount of resources (or energy) remains constant, framing evolution as a zero-sum game. In this paradigm, any evolutionary advantage gained by one lineage is offset by an equivalent disadvantage for a potential competitor (or predator) within the same adaptive zone (Van Valen, 1973; also discussed in Stenseth & Smith, 1984). Therefore, evolution as proposed by Van Valen is a memoryless process and age should not influence a given lineage’s probability of extinction.

After Van Valen’s seminal work, very few studies have been conducted on this specific topic, most of them at higher taxonomic levels than the species. The vast majority of those that have examined clades at the species level have demonstrated age-dependent extinction (Januario & Quental, 2021, and references therein). These examples present either positive age dependency — an increase in the probability of extinction linked to age (Pearson, 1995; Parker & Arnold, 1997; Doran *et al*., 2006; Ezard *et al*., 2011) or negative age dependency —a decrease in the probability of extinction linked to age (Jones & Nicol 1986; Finnegan *et al*., 2008; Crampton *et al*., 2016; Hagen *et al*., 2018; Silvestro *et al*., 2020). Hypothesis behind negative age-dependent extinctions typically evokes stochastic demographic effects associated with small population size and small geographical range, which result in a higher extinction probability for younger species. (Liow & Stenseth, 2007; Miller, 1997; Payne & Finnegan, 2007; Foote *et al*., 2008; Jablonski, 2008). Those entailing positive age dependency range from evolutionary ratchet mechanism that causes “species senescence” and reduced evolvability (akin to the process discussed by Muller, 1964), to macroevolutionary ratchet mechanisms resulting in increased specialization (Van Valkenburg, 1999; Valkenburgh *et al.,* 2004), to a scenario of morpho-ecological stasis, where old species may not be able to compete against new species (Pearson, 1995; Eldredge *et al.,* 2005).

The discrepancy observed between Van Valen’s original work and subsequent studies can be understood by considering the different taxonomic levels employed in these investigations (Januário & Quental, 2021). In his initial research, Van Valen (1973) assumed that higher taxonomic categories could serve as proxies for understanding dynamics at the species level. However, among the 25,000 sub-taxa distributed across 46 distinct clades, only four clades were studied at the species level. It’s worth noting that only a limited number of studies have explored the implications of using various taxonomic levels when inferring age-dependent extinction patterns. Pearson (1995) found positive age dependency in both species and genera of trilobites, Ezard *et al*. (2016) positive age dependency at the species level and age-independent extinction at the genus level (planktonic foraminifera), and Januario & Quental (2021), examining fossils of ruminants, negative age dependency at the species level, and age independence at the genus level.

Additionally, the vast majority of previous studies have examined the possibility of age-dependent extinction by defining their study groups either phylogenetically or taxonomically (e.g., Pearson, 1995; Crampton *et al*., 2016; Condamine *et al*., 2019; Silvestro *et al*., 2020). Hence most studies do not explicitly incorporate ecology (but see Ezard *et al.,* 2011), except for assuming that species within each clade share ecological similarities (which may be a reasonable assumption for many groups). The difficulty of using ecology to identify the pool of species belonging to a given adaptive zone is echoed in Van Valen’s work (1973), where the challenge of defining these homogeneous groups is briefly discussed, but higher taxa were used as a convenient approximation for such adaptive zones. Although subsequent work continued to use a taxonomically or phylogenetically defined pool of species when testing the “Law of Constant Extinction” (e.g., Pearson, 1995; Crampton *et al*., 2016; Condamine *et al*., 2019; Silvestro *et al*., 2020), an ecologically defined pool of species would more closely align with Van Valen’s original idea of the adaptive zone and his underlying Red Queen hypothesis. Additionally, differences in ecology (or morphological traits used as a proxy for ecology) have been shown to affect extinction probabilities (Van Valkenburgh 1999; Chichorro *et al*., 2019; Hembry & Weber, 2020; Zeng & Wiens 2021), implying that considering such effects when attempting to examine age-dependent extinction probability could be an important step toward gaining a deeper understanding of extinction risk.

Here we used the North American Canidae fossil record to investigate how different pools of species (i.e. phylogenetic pool and ecological pool) affect our capacity to understand the “Law of Constant Extinction”. The distinct phylogenetic pools of species were based on well-defined taxonomy and phylogenetic reconstruction of the group (Wang & Tedford, 2008; Slater, 2015). Specifically, we delineated the entire Canidae family and its three subfamilies—Hesperocyoninae, Borophaginae, and Caninae as different phylogenetic defined pools of species. The ecological pools of species were defined according to distinct dietary guilds described for the Canidae family (Van Valkenburgh, 1991; Wang & Tedford, 2008; Slater, 2015), namely, hypercarnivores (more than 70% of its diet consists of large-vertebrates preys), mesocarnivores (a diet of about 50 to 70% of vertebrate material), and hypocarnivores (small predators that use invertebrates and plant material and consume less than 30% of vertebrate material). Defining species based on guilds not only offers a representation of an adaptive zone more similar to Van Valen’s original work but also takes into account the fact that these ecological guilds have demonstrated significant macroevolutionary consequences (Van Valkenburgh & Damuth, 2004; Balisi *et al*., 2018). More specifically we tested the following hypothesis: 1-The signal of age-independency is more evident in the ecological pool of species than in a phylogenetic pool of species; 2-When compared to other dietary guilds, hypercarnivores would show a stronger tendency for negative age-dependent extinction. The first hypothesis was based on the idea that the ecological pool of species provides a more accurate representation of the adaptive zone used by Van Valen (1973), where the Red Queen would act as the primary force steering age-independent extinction. The second hypothesis was based on previous work that showed that: 1- species with smaller body size have greater population density relative to those of larger species (Damuth, 1981; Damuth, 1987); 2- higher population density lowers the probability of extinction (Purvis *et al*, 2000); 3- the level of carnivory is related to body size (Valkenburgh *et al*., 2004; but see Balisi & Van Valkenburg, 2020); 4- large body size is related to small litter and slow reproduction rates which could enhanced extinction probability (McKinney, 1997; Purvis *et al*., 2000). We note however that large hypercarnivores also have large geographical distributions which typically decrease the probability of extinction (Payne & Finnegan, 2007; Foote *et al*., 2008; Jablonski, 2008).

## Methods

### Study System, Ecomorphological data, and Phylogenetic imputation

The Canidae family has a well-documented fossil record both in space and time (Wang, 1994; Wang *et al*., 1999; Tedford *et al*., 2009). There are thorough ecomorphological characterizations available for the majority of both extinct and extant species (Janis *et al*., 1998; Van Valkenburgh *et al*., 2004; Slater, 2015; Balisi *et al*., 2018; Balisi & Van Valkenburgh, 2020) and the group shows a diverse spectrum of body sizes and dietary preferences. The family’s evolutionary history is largely centered in North America (Wang & Tedford, 2008), the continental region studied here, making it a study system with a well-defined geographical arena. We downloaded and curated data on body mass and craniodental variables from different sources. Body mass information and measurements of the lower first molar (m1) for 140 canid species were taken from the dataset published by Faurby *et al*. (2021), which compiles information from various literature sources. Craniodental measurements, including relative blade length of the lower first molar (RBL), lower first molar blade size relative to dentary (lower jaw) length (M1BS), the size of the lower second molar relative to the dentary (lower jaw) length (M2S), the mechanical advantage of the temporalis muscle (MAT), the robustness of the lower fourth premolar (p4S), and relative lower molar grinding area (RLGA) were taken from the dataset published by Slater (2015). Following the approach described by Slater (2015), these craniodental variables were used to characterize the diet of all fossil species (see analysis details below and Supplementary material) into three dietary categories (hypercarnivores, mesocarnivores, and hypocarnivores). These variables have been suggested as reliable indicators of dietary groups in canids (Van Valkenburg & Koepfili, 1993). Due to the nature of fossil material, our dataset exhibited varying degrees of missing data (Table S1), ranging from 1.5% to 51%, with an average of 32% of missing data. To address the issue of missing data, we applied a phylogenetically informed imputation approach, implemented in the R package *Rphylopars* (Goolsby *et al*., 2017). The final morphological dataset included observed and imputed values for the 6 craniodental variables that showed strong phylogenetic signal (Fig. S1) for 142 canid species, as well as observed values for the MAT variable for 69 extant and fossil species (for more details about the imputation process see Supplementary Material). While we did not impute the MAT variable due to its low phylogenetic signal and high degree of missing data, it was utilized in the ecological characterization of 69 fossil species. To evaluate the accuracy of the estimated values for craniodental variables, we created a simulated dataset with a proportion of missing data similar to that of the original dataset for species with complete data. We then compared the imputed estimates with the observed empirical values. The results demonstrate a strong positive correlation between the empirical data and the imputed estimates, confirming the reliability of the imputation method (Fig. S2).

### Ecological categorization using discriminant analysis

Following Slater (2015), we used a Linear Discriminant Analysis (LDA) to characterize the diet of fossil species into one of three dietary categories (hypercarnivore, mesocarnivore, or hypocarnivore). The LDA was implemented in the *Applied Statistics with S* (MASS) (Venables & Ripley, 2002) package for R. The scripts used here were strongly based on the scripts provided by Slater (2015). In the LDA analysis, we used a training set (based on the training set used by Slater (2015)), encompassing extant canids and other carnivores, summing up 35 species. The training set was composed of 25 extant canid species, nine procyonids, and *Ailurus fulgens*. The species selected for the training sets were chosen based on their demonstrated effectiveness in the diet classification of fossil species (Slater, 2015), and data availability. We used a diet classification for the training set proposed by Hopkins *et al*. (2021) with 4 hypercarnivores, 17 mesocarnivores, and 14 hypocarnivores. Using a diet classification proposed by Slater 2015, results in very similar inferences (see supplemental material) hence we kept the most recent classification by Hopkins *et al*. (2021).

We conducted two separate discriminant analyses since we were unable to impute MAT variable. In the first LDA, we used the training sets with all six variables to establish the association between the craniodental variables and the different diet categories, generating a multivariate linear model. We employed the trained LDA model to generate predictions of the dietary categories of the extinct canids. We successfully characterized 69 fossil species. In the second LDA, we excluded the MAT variable (because several fossil species do not have this variable) and followed the same procedure. Due to the inference of very few hypocarnivores species and considering that such discrimination procedure is more effective at separating hypercarnivores from the other categories (Van Valkenburgh & Koepfli, 1993; Slater, 2015), we decided to merge mesocarnivores and hypocarnivores, in all subsequent analyses, as unique category “non-hypercarnivores”. Out of our 142 species, only two – *Canis feneus* and *Cynarctoides roii* – could not be classified using our approach due to a lack of morphological data and their absence from the phylogeny. To include these species, we adopted the classifications proposed by Balisi & Valkenburg (2020), which relied on the length of the blade on the lower molar (carnassial) relative to dentary length (m1BS).Consequently, our final ecological classification comprised 44 hypercarnivores and 98 non-hypercarnivores.

### Fossil occurrence data

We used two different fossil occurrence datasets. The first one (hereafter dataset 01) was compiled and curated by Lucas M. V. Porto (Porto, L.M. V., personal communication, 2023) for a taxonomic broader-scale project. The fossil occurrences from the North American and Eurasian Carnivora dataset were compiled from online databases: Paleobiology Database (PBDB; https://paleobiodb.org/#/) and New and Old Worlds fossil mammal database (NOW; https://nowdatabase.org). The curatorial work carried out adopted a conservative measure regarding species taxonomic uncertainty. It kept only described species and filtered out all occurrences without or with uncertain species identification, removing all occurrences that were tagged with taxonomic uncertainty markers (Bengtson, 1988; Sigovini *et al*., 2016): “indet.”, “Sp.”, “?”, “Aff.”, “Cf.”. Faurby *et al*. (2019) were used to recombine subjective synonyms and subspecies. To remove occurrences with lower temporal resolution, all occurrences with an estimated time interval of more than 6 million years were removed. Given the fossil dataset used is compiled from two distinct databases, it is important to acknowledge the potential presence of duplicates. In this case, duplicates refer to situations where data points are mistakenly treated as independent replicates when they are the same occurrence. This can occur due to overlap or redundancy in the information provided by the databases. It can lead to an overestimation of the sample size and potentially introduce bias into the results. To address the issue of duplication (“pseudoreplication”) only fossil occurrences with a temporal range that differed from each other by at least 6 million years and were spatially distant by at least 1 degree in latitudinal and longitudinal coordinates were retained. After all the curatorial work, the final data set was left with 1735 occurrences of 133 species. The occurrences for North American Canidae were kindly supplied by Lucas Porto for this project. The second fossil occurrence dataset (hereafter dataset 02) was taken from Balisi & Valkenburgh (2020). This dataset uses occurrences from PBDB and the Neogene Mammal Mapping Portal (NEOMAP, http://ucmp.berkeley.edu/neomap). This database encompasses more occurrences and an average better temporal resolution since Balisi & Valkenburgh (2020) data is mostly composed of data derived from NEOMAP. This dataset was used to compare the results derived solely from the PBDB. This dataset compiles 3708 fossil occurrences (most from a historical temporal scale of less than 0.01 Ma) of 138 species.

### Initial diversification analyses

To examine the impact of age on the probability of extinction, it is better to conduct diversification analyses within a time window where the background extinction rate remains constant (Hagen *et al*., 2018). This is crucial because the lifespan of species is inversely proportional to extinction rates (Raup, 1985). Therefore, changes in the background extinction rates would influence lineage duration and interfere with the estimation of age dependency. To establish the appropriate time windows for subsequent analysis, we initially inferred the temporal variation in speciation and extinction rates. For the diversification analysis, we examined, in both fossil occurrences datasets, the speciation and extinction dynamics for 6 distinct species pools. These included pools defined by dietary classification, such as Hypercarnivores and Non-hypercarnivores (ecological pool of species), as well as those defined phylogenetically, such as the entire Canidae family and its three sub-families: Hesperocyoninae, Borophaginae, and Caninae (phylogenetic pool of species). All analyses were conducted at the species level.

We employed a birth-death model approach implemented within a Bayesian framework using the software PyRate (Silvestro *et al*., 2014, 2019) to estimate the diversification rate over time. This model simultaneously estimates; 1- the times of origin and extinction; 2- rates of speciation and extinction; 3- if those rates change through time; and 4- the preservation rate, while taking into account several aspects of the incompleteness of the fossil record. To account for age uncertainties in our analyses of dataset 01, we generated 50 replicated resampled datasets for each species pool by randomly sampling a point time within the stratigraphic interval of each occurrence, resulting in a total of 300 replicated datasets (50 replicates * 6 different datasets, each representing a different pool of species). Each of these 300 replicated datasets was used in the diversification analysis to describe the temporal trend of diversification rates. Prior to the diversification analysis, we ran a maximum likelihood test to assess which preservation model is best supported by one of those datasets (Silvestro *et al*., 2019) (Supplementary Material). We used the best preservations model for each analysis coupled with a Gamma model, which allowed for preservation rate heterogeneity across lineages. We used a reversible jump Markov Chain Monte Carlo (RJMCMC) to jointly estimate the model parameters, including the number and temporal placement of rate shifts in speciation and extinction (Silvestro *et al*., 2019). We ran 60 million iterations and sampled at every 10,000 to achieve convergence. For dataset 02, we followed the same steps, but instead of 50 replicates, we utilized 25, yielding a total of 150 replicate datasets (25 * 6). The Bayes Factor (Kass & Raftery, 1995) was used to test the evidence in favor of a given temporal shift in diversification rates (speciation or extinction). The effective sample sizes (ESS) were assessed by visualizing the log files in Tracer (Rambaut *et al.,* 2018) after excluding the first 10% of the sample as a burn-in period. All replicates had sufficient sampling (ESS values > 200).

Based on the diversification analysis results, we identified different time windows across all species pools where the background extinction rate remained constant. Notably, the diversification dynamics observed in dataset 02 (Fig. 1-2 and Fig. S4-5) present some small differences (e.g. the exact position of a given shift; the presence or absence of a small shift) from those in dataset 01, but as we will see do not result in different inferences down the line. As dataset 01 serves as our primary analysis, we will only present the time windows described for this dataset here but see the Supplementary Material for an overview of all analysis. Our diversification analyses revealed four distinct time periods within the ecological species pool (Fig. 1): one for hypercarnivores (Hypercarnivores 01), from the root age (defined by a 95% HPD between 35.6 and 32.6 mya) to 6.5 mya (Fig. 1A). Three for non-hypercarnivores (Non-hypercarnivores 01), spanning from the root age (defined by a 95% HPD between 39 and 37.3 mya) to 18 mya, Non-hypercarnivores 02, from 14.5 mya to 2 mya, and Non-hypercarnivores 03, from 14.5 mya to the present (Fig 1B).

**Figure 1.**
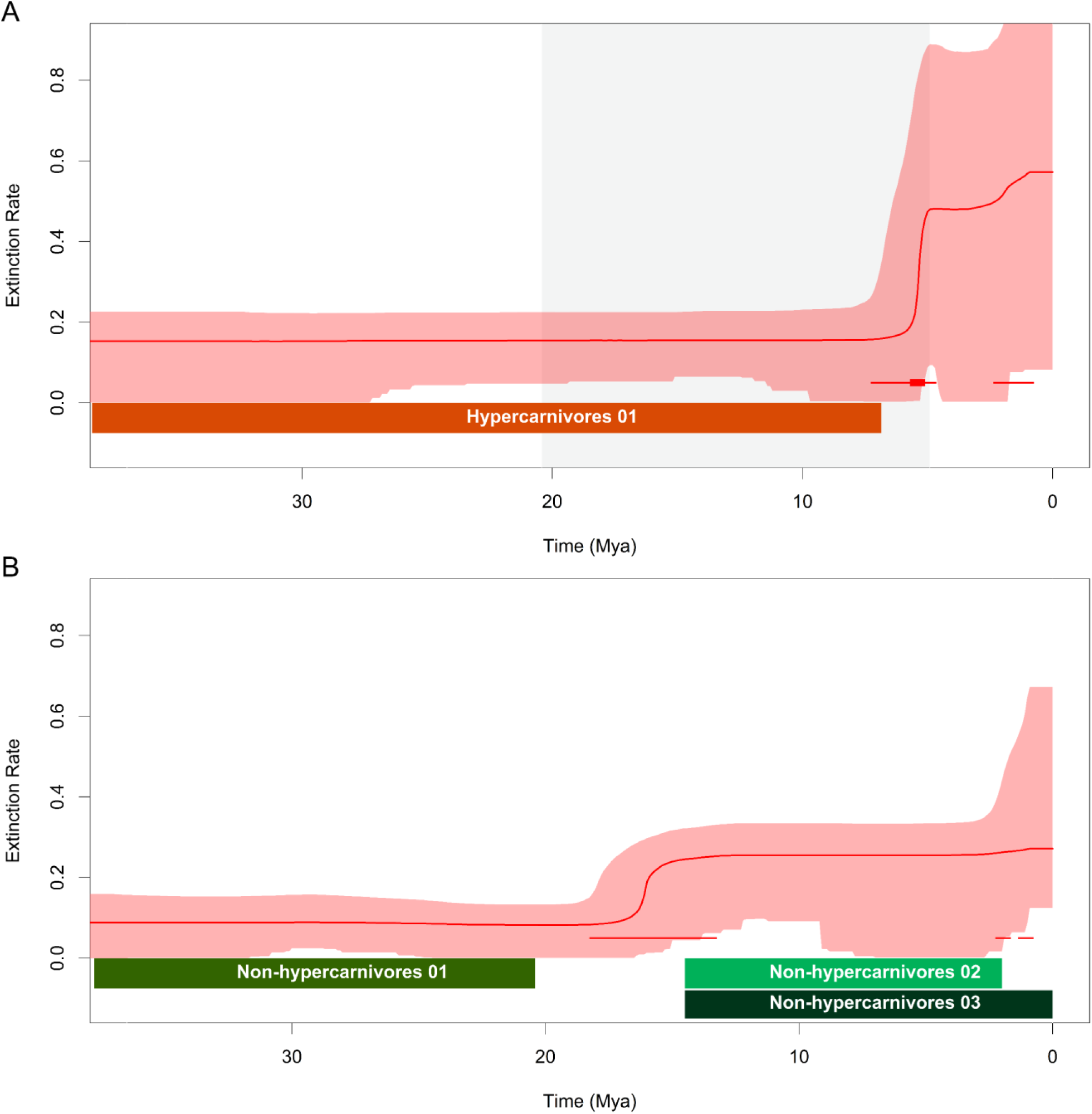
Extinction rate for hypercarnivores and non-hypercarnivores for the first dataset (PBDB). Panels A and B display plots of extinction rates over time, with the light-colored area representing the 95% highest posterior density interval (HPD) for the extinction rate of hypercarnivores (Panel A) and non-hypercarnivores (Panel B). The solid line within the light-colored area represents the mean of the posterior distribution of rate values at each time point. Red horizontal segments indicate times of low significance (2 < BF < 6) rate shifts in extinction, while red rectangles mark times of highly significant (BF > 6) rate shifts in speciation and extinction. In panel A, the white and grey background bars indicate the preservation intervals used in the analysis, allowing preservation to vary among but remain constant within intervals (preservation variation among lineages was allowed within each time window). Colored horizontal bars indicate the length of the different time windows used in the ADE analysis. Windows with shifts in the extinction rate were included to examine the effect of changes in extinction rate on the inference of age-dependent extinction.

**Figure 2.**
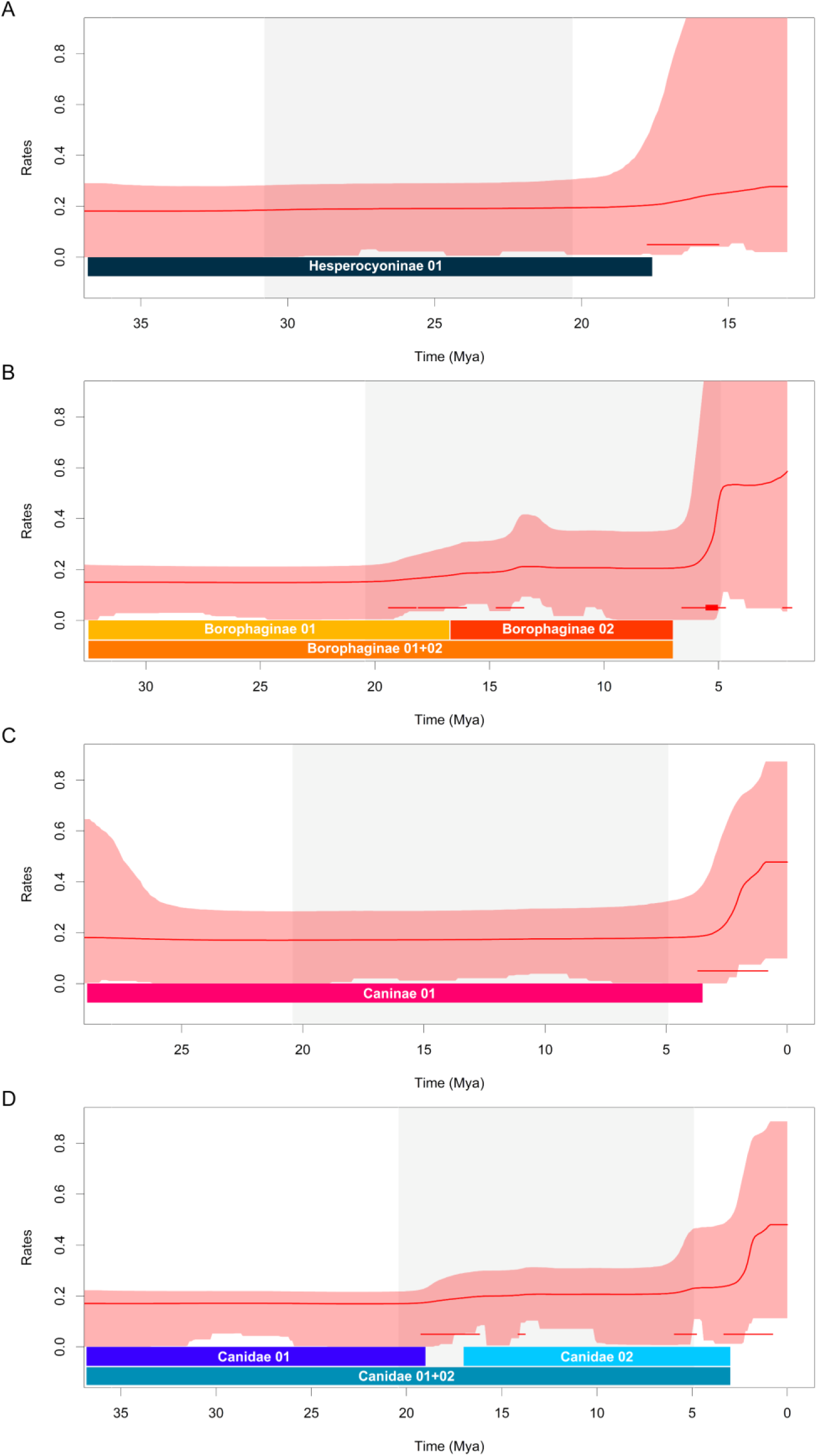
Extinction rate through time for Canidae and its subfamilies. Panels A, B, C, and D show extinction rate through time, with the light-colored area representing the 95% highest posterior density interval (HPD) for extinction rate for Hesperocyoninae (Panel A), Borophaginae (Panel B), Caninae (Panel C), and Canidae (Panel D). The continuous line inside the light-colored area indicates the mean of the posterior distribution of rate values at each moment in time. Red horizontal segments signify times of low significance (2 < BF < 6) rate shifts in speciation and extinction rates, while red rectangles represent the times of highly significant (BF > 6) rate shifts in extinction. The white and grey background bars indicate the preservation intervals used in the analysis where we allowed preservation to vary among but be constant within intervals (preservation variation among lineages was allowed within each time window). The horizontal-colored bars on the bottom of panels (A), (B), (C), and (D), indicate the length of the different time windows used in each ADE analysis. Please note: the windows that encompass shifts in the extinction rate (Borophaginae 01+02 and Canidae 01+02) were added to investigate how the shift would impact the inference of age dependency. Windows with shifts in the extinction rate were included to examine the effect of changes in extinction rate on the inference of age-dependent extinction.

Regarding the phylogenetic species pool, we identified eight distinct time periods (Fig. 2). One for Hesperocyoninae (Hesperocyoninae 01), from the root age (defined by a 95% HPD between 37.4 and 36.8 mya) to 17.6 mya (Fig. 2A). Three for Borophaginae (Borophaginae 01), from the root age (defined by a 95% HPD between 35.6 and 33.4 mya) to 16.75 mya, Borophaginae 02, from 16.75 mya to 7 mya, and Borophaginae 01+02, from the root age (defined by a 95% HPD between 35.6 and 33.4 million years ago) to 7 mya (Fig. 2B). One for Caninae (Caninae 01), from the root age (defined by a 95% HPD between 32.2 and 28.6 mya) to 3.5 mya (Fig. 2C). Three for Canidae (Canidae 01), from the root age (defined by a 95% HPD between 37.5 and 36.9 mya) to 19 mya, Canidae 02, from 17 mya to 3 mya, and Canidae 01+02, from the root age to 3 mya (Fig. 2). We delineated some windows with shifts in the extinction rate (Fig. 1B – Non-hypercarnivores 03; Fig. 2B – Borophaginae 01+02; Fig. 2D – Canidae 01+02) to examine the impact of these minor shifts on the inference of age-independent extinction. For more detailed information on how the time periods were defined, please refer to the Supplementary Material

### Age-dependency analyses

To infer the degree of age-dependency (or independence) on extinction rate dynamics we followed the framework described by Hagen *et al*. (2018) and implemented it in PyRate. In the age-dependent extinction model (Hagen *et al.,* 2018), lineage duration is modeled according to a Weibull distribution where the shape parameter describes the age-dependence effect on extinction. For a shape parameter > 1, the probability of extinction increases with the age of the taxon; for a shape parameter < 1, the extinction is greater for younger lineages, and it declines with age (Fig. S3). If the shape parameter is equal to or close to 1, then extinction is interpreted to be age-independent, being congruent with the “law of constant extinction” described by Van Valen (1973). In this case, the Weibull distribution reduces to an exponential distribution, which characterizes a simple birth-death model. This framework also explicitly takes into account species that have not been sampled (Hagen *et al.,* 2018). Those are typically short-lived (Foote & Raup, 1996) and their absence could strongly influence our inference of the shape parameter, and hence extinction age-dependency. In this Bayesian framework, fossil occurrences are used to simultaneously estimate: (1) the “true” times of origination/speciation and extinction of each sampled lineage; (2) the preservation rate for each predefined time window; (3) the parameters of the modified Weibull, including the shape parameter that measures the association between the probability of extinction and the duration of the species.

We ran the age-dependent analysis in each one of the different time windows shown in Figs. 1-2 and Fig. S4-5. In each window, we ran a maximum likelihood test to determine which preservation model is best supported by the data (Table S2 and Table S3). The age-dependent analysis only allows for the use of the homogeneous Poisson process (HPP) and the time-dependent Poisson process (TPP) models. Therefore, if NHPP was the best model, we selected the second-best model between HPP and TPP. The timespan and the chosen preservation model are summarized in Table S2-3. The Gamma model was not employed in the age-dependent analysis as it has not been fully tested and preliminary tests showed that in our empirical case, it mistreated singletons when applying the age-dependent model. We ran the RJMCMC integrator setting the age-dependency model described for each one of the different time windows described above using 20 million iterations sampled every 10.000 iterations to obtain the posteriors for each parameter. The ESS was assessed as previously described. We compiled the results for all the replicas within a single posterior distribution to summarize the parameter estimation (shape) of the Weibull-modeled distribution of lineage durations in the different species pools. To evaluate the evidence in favor of age-dependent extinction, we compared the 95% highest posterior density (HPD) of the shape parameter of each species pool against the reference value of 1 which defines an age-independent extinction.

## Results

### Age-dependent analyses

Although the time windows in dataset 01 and dataset 02 slightly differed both in their number and duration, the ADE findings were remarkably similar (Fig. 3; Fig. S7). Therefore, we chose to present here only the results derived from dataset 01 (see supplementary material for all results). In our analysis of the ecological species pool, we observed distinct dynamics between hypercarnivores and non-hypercarnivores (Fig. 3). Hypercarnivores displayed a pattern more consistent with positive age-dependent extinction, while non-hypercarnivores leaned towards age-independent extinction. It is worth noticing that in the window with a small extinction shift the ADE signal changes and in this particular case presented strong evidence for negative age-dependent extinction. Therefore, different ecological pools of species showed varied signals of age dependency or independence.

**Figure 3.**
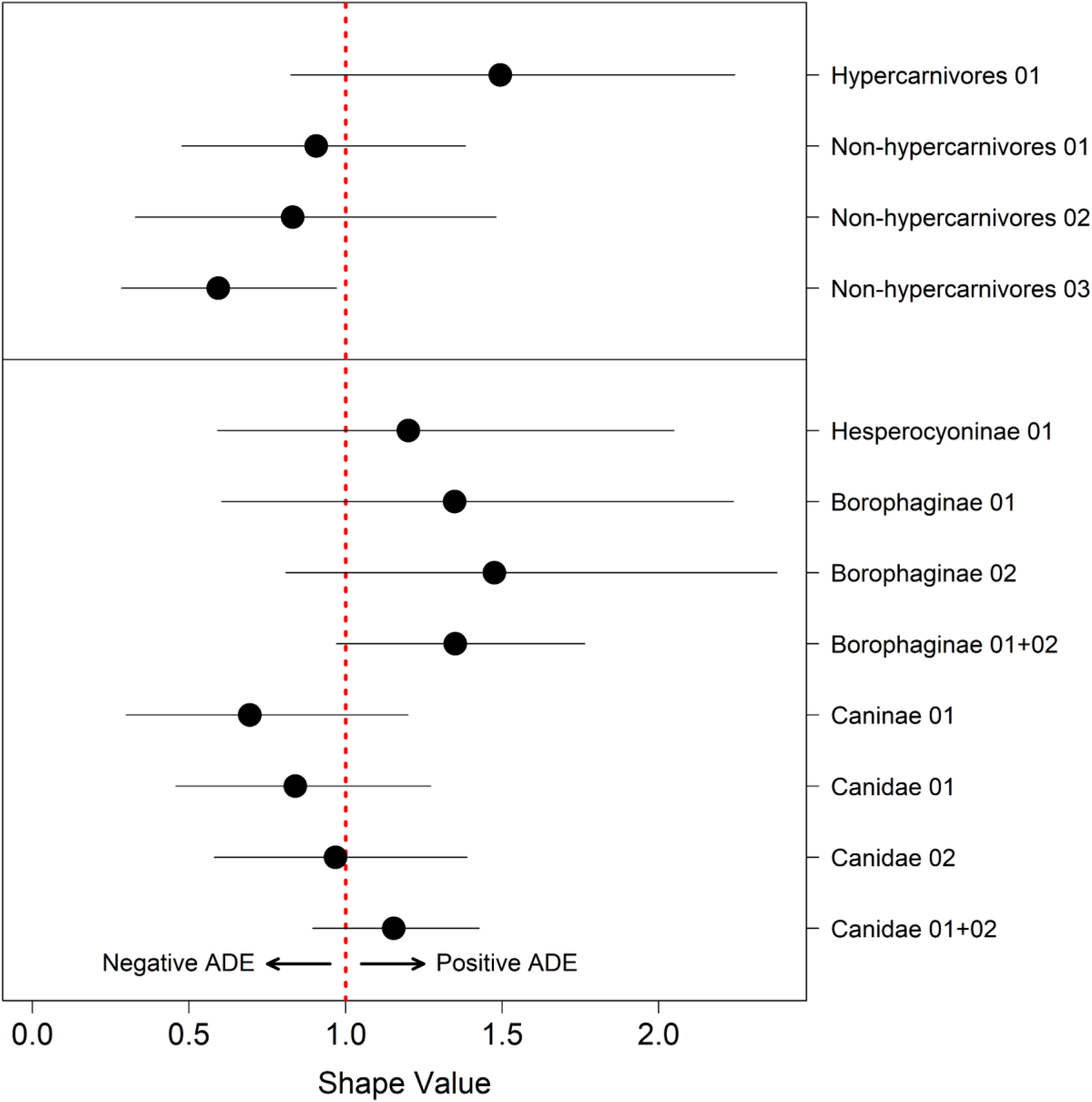
Extinction age-dependency for the ecological pool of species (hypercarnivores and non-hypercarnivores) and the phylogenetic pool of species (Canidae and its subfamilies). Posterior distribution for the shape parameter of the Weibull distribution for each time window. The horizontal black lines indicate the 95% highest posterior density (HPD) interval, and the black dots represent the median of each posterior distribution The red vertical line indicates age-independency (shape value = 1).

In hypercarnivores, we found moderate evidence in favor of a positive age-dependent extinction at the Hypercarnivore 01 window. Although the 95% HPD (Highest Posterior Density) interval estimate for the shape parameter (median = 1.5, 95% HPD = 0.79 – 2.23) includes the value of 1 (Fig. 3), there is a suggestive trend towards positive age-dependent extinction. When visually inspecting the shape estimates for each replicated dataset individually (Fig. S8A), we observe that all replicates show their posterior shifted toward values higher than 1, with a few replicated datasets providing strong evidence for positive age dependency. This consistent pattern persists across all three windows examined in dataset 02 (Fig. S6B).

In non-hypercarnivores, we reject age dependence as we found evidence for age-independence in two windows analyzed. In the Non-hypercarnivores windows 01 and 02 (Fig. 3), the median of the posterior distribution of the shape parameter is very close to 1, particularly in window 01 (window 01: median = 0.90, 95% HPD = 0.48 – 1.39; window 02: median = 0.83, 95% HPD = 0.33 – 1.49). In both windows, the 95% HPD is reasonably centered around the value of 1 but in Non-hypercarnivores 02 window (Fig. 3) the posterior is slightly biased toward values smaller than one. By visually examining the estimated shape values for each replicated dataset individually within both windows we found a similar pattern (Fig. S8B – C). Several of the individual replicated datasets demonstrate a slightly biased toward values smaller than and several replicated datasets providing strong evidence for an age-independence extinction dynamic. Similarly, results are seen for dataset 02, with the Non-hypercarnivores 01 window presenting even stronger evidence for age-independent extinction (Fig. S7B). When a small shift in background extinction is ignored and a longer window is analyzed (Non-hypercarnivores 03 window), we observed a clear signal for negative age-dependency (Fig. 3). The estimated shape parameter of the Weibull distribution (median = 0.59, 95% HPD = 0.29 – 0.96) was significantly smaller than 1, providing strong evidence for negative age dependency. Examining the estimated shape values for each replicated dataset we can see that the majority of replicated dataset provide strong support for a negative age-dependency (Fig. S8D).

Our analysis of the phylogenetic species pool, which includes Canidae, Hesperocyoninae, Borophaginae, and Caninae, suggests a scenario more in accordance with age-independency extinction for most time windows (Fig. 3), but there is some considerable variation among the different windows. In Hesperocyoninae 01 (Fig. 3), we observe a wide posterior distribution, with a positive median close to 1 (median = 1.19, 95% HPD = 0.55 – 1.97). Upon visually inspecting the replicated datasets, we notice that several replicated datasets suggest strong evidence for age independence since most posterior estimates are centered very close to 1, while others indicate weak evidence in favor of positive age dependency (Fig. S9A). In the three windows of Borophaginae, we notice some tendency towards positive age-dependent extinction (Fig. 3): Borophaginae 01 (median = 1.34, 95% HPD = 0.63 – 2.24), Borophaginae 02 (median = 1.47, 95% HPD = 0.75 – 2.31), and Borophaginae 01+02 (median = 1.35, 95% HPD = 0.97 – 1.76). When we visually inspect the replicated datasets the individual replicates for Borophaginae 01 and 02 show more variation than the Borophaginae 01+02 window, where each replicate dataset consistently shows a stronger tendency towards positive age dependency (Fig. S9B – D). We note that Borophaginae 01+02 combines the other two windows and includes a rate shift in extinction, which could impact the estimates of the shape parameter. Conversely, Caninae 01 (median = 0.70, 95% HPD = 0.30 – 1.20), exhibited a weak tendency towards negative age-dependent extinction (Fig. 3). Upon visual inspection of the replicated datasets, we can see that all replicates show their posterior distributions shifted towards values lower than 1, with some replicated datasets providing evidence for negative age-dependency (Fig. S9E). In Canidae (Fig. 3), the shape parameter for each window was found to be very close to 1: Canidae 01 (median = 0.84, 95% HPD = 0.45 – 1.24), Canidae 02 (median = 0.96, 95% HPD = 0.59 – 1.38), and Canidae 03 (median = 1.15, 95% HPD = 0.89 – 1.44). By visually examining the replicated datasets of Canidae 01 and Canidae 02, we notice that in several replicated datasets, most posterior estimates are centered close to 1, which supports age independence (Fig S9F – G). In Canidae 01+02, we found variable support for a positive age dependency in some replicates, although all show a median above 1 (Fig. S9H). We note this time window encompasses a rate shift in extinction as in Borophaginae 01+02

To help visualize the interplay between different ways of creating the species pool (by diet or phylogeny), and the effect of missing species, we plotted the empirical longevities alongside the estimated Weibull curves (red lines in Fig. 4 and Fig. S10). These curves were constructed using the shape and scale parameters derived from PyRate’s age-dependent model and superimposed over the empirical distributions of species longevities, which were color-coded according to their diet or phylogenetic affiliation (histograms in Fig. 4; see also the supplemental material for extra panels). The dark blue line indicates the expected distribution of lineage duration under age-independent extinction. The visualization suggests that the estimated curves fit the empirical longevity distributions reasonably well, although for some windows the preservation model suggests there is a reasonable proportion of species with short longevities missing from the empirical distribution due to preservation biases.

**Figure 4.**
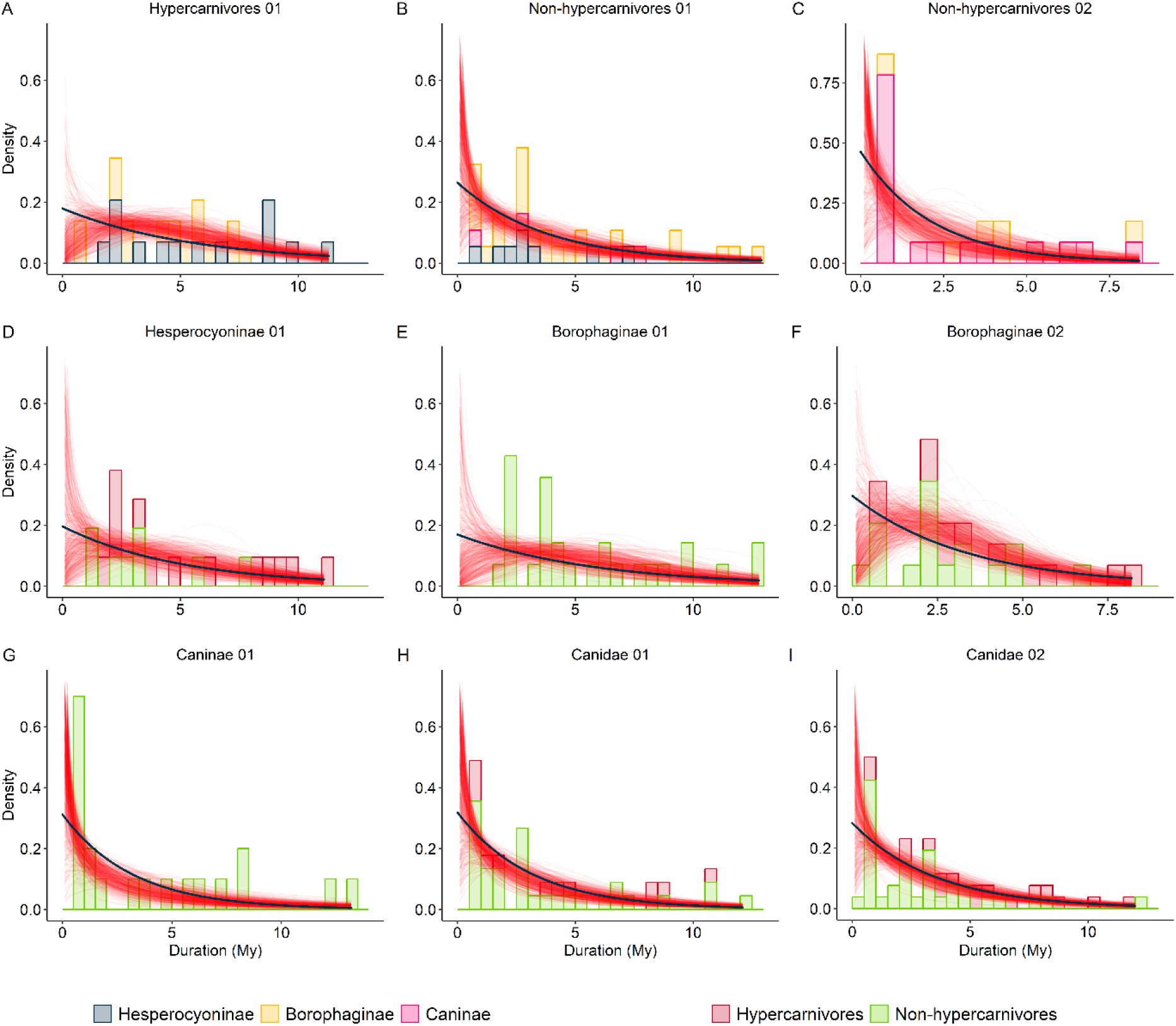
Empirical distribution of species durations estimated in PyRate for the sampled taxa, and the results for the age-dependent model at each window. Empirical longevity estimates were taken for the combined posterior distribution using all replicates by taking the difference between the estimated median values for the times of speciation and extinction. In all panels, the color bars indicate the diet categories of Canidae (Red: hypercarnivores, and Green: non-hypercarnivores) or the sub-families (Grey: Hesperocyoninae, Yellow: Borophaginae, Pink: Caninae). Within each panel, the Weibull distributions estimated for different iterations by PyRate’s age-dependent model are depicted using 500 semitransparent red lines, randomly drawn from the combined posterior. Expected distribution of lineage duration under an age-independent extinction is shown in the dark blue line.

Interestingly, the empirical longevity distribution of windows of the phylogenetic pool of species showing a tendency for positive age-dependent extinction resembles that of hypercarnivores. With the exception of Borophaginae 01 (see discussion), those windows show a higher proportion of hypercarnivore species within it. On the other hand, the empirical longevity distribution of windows of the phylogenetic pool of species showing a tendency for negative age-dependent extinction resembles that of non-hypercarnivores and shows a very low, or even zero, proportion of hypercarnivores. Within the studied time window, the hypercarnivore pool consists solely of Hesperocyoninae and Borophaginae species. In contrast, non-hypercarnivores exhibit distinct patterns (Fig. 4). For Non-hypercarnivores 01 (Fig. 4B), there are species from all sub-families, although a preponderance of Hesperocyoninae and Borophaginae species. For Non-hypercarnivores 02, we only see Borophaginae and Caninae species, with strong preponderance from the latter.

## Discussion

Here we investigated how different species pools defined by ecology (diet categories) or phylogeny/taxonomy interfere with our inference of the “Law of Constant Extinction”. We focus our discussion on results from windows without shifts in the extinction rate but return to those including shifts when discussing methodological implications. We found different signals of age-dependent extinction when using different species pools defined by phylogeny or ecology (Fig. 3). When using the phylogenetic species pool, our results revealed a broader spectrum of signals concerning age dependency or age-independency in extinction patterns. Some analyses have provided compelling evidence for negative age-dependent extinction (notably observed in Caninae 01 and Canidae 01), others have pointed to weaker evidence in favor of a positive age-dependent extinction (as evident in Borophaginae 01, Borophaginae 02), and there are instances where evidence strongly supports age-independent extinction as is the case of Canidae 02. The ecological pool of species either supports age-independency (Non-hypercarnivores) or positive age-dependency (Hypercarnivores).

When looking at each sub-family separately we noticed opposing signals. Borophaginae and Hesperocyoninae showed positive shape values and Caninae showed negative ones. The result for the whole family showed a signal more akin to age-independent extinction, in particular the Canidae 02 window, inviting one to wonder if this signal at the family level emerges as an “average” result from combining the three sub-families. In a previous study on carnivores, a negative age-dependent extinction was identified at the Order level (Hagel *et al*., 2018). Hence the age-dependent signal might shift from positive (Borophaginae 02), to age-independent (Canidae 02), to negative (whole Carnivora – Hagen *et al*., 2018) depending on the taxonomic rank used to delimit the species pool. Additionally, we might or not observe changes in the age-dependent signal for the same taxonomic pool of species at different time windows. Within the Canidae family, we observed distinct signals of age-dependency across different time intervals, aligning with similar observations made by Crampton *et al*. (2016), who showed that the signal of age-dependent extinction might change over time. It is also worth noting that the relative composition of the three sub-families changed as time went by, reinforcing the idea that the signal recovered at a higher phylogenetic level results from the combination from combining their sub-clades. The variability in the signals of age-dependency across different taxonomic ranks and among different time windows confirms the expectation that the pool of species strongly influences the age-dependent signal of extinction and invites one to think about what those differences mean and how the ecological composition within each phylogenetically defined pool of species might interfere with our inference.

Even though hypercarnivory evolves in all three Canidae sub-families, it is more predominant in the Hesperocyoninae (17 hypecarnivores out of 27 species) and Borophaginae (21 hypecarnivores out of 70 species) subfamilies than in Caninae (6 hypercanivorous species out of 45 species). This discrepancy is reflected in our sampling analysis. In our time windows we have: 0 hyper out of 28 in the time window Borophaginae 01, 12 hyper out of 29 in the time window Borophaginae 02, 13 hyper out of 21 in the time window Hesperocyoninae 01, and not a single hypercarnivore in time window Caninae 01. Although not perfect (but see discussion below), we see a tendency for those phylogenetic species pools that contain more hypercarnivores to be more likely to present positive-age dependency. Notably, in the ecological species pool, our findings revealed evidence of positive age-dependent extinction among hypercarnivores and age-independent extinction among non-hypercarnivores (same for dataset 02; see Fig. S7B). We argue that the age-dependent extinction signal observed when using the ecological pool of species helps us explain the different signals we found when using the phylogenetic.

Sub-families with a higher proportion of hypercarnivores (Borophaginae and Hesperocyoninae) displayed more positive estimates of the shape parameter compared to the Caninae subfamily, which in our time window is absent of hypercarnivores and revealed a slightly negative shape parameter estimate. We also note that the Borophaginae 02 window showed a much stronger signal of positive age-dependent extinction and a higher proportion of hypercarnivores in that pool, in accordance with the ecological species pool reasoning. However, one notable exception is the result for Borophaginae in window 01, which showed a significantly positive shape parameter value, despite the absence of any hypercarnivore species in that time window. The 28 species within that window exhibit significant differences in both body mass and LD1 values (a proxy for carnivory) when compared to other canids (see Fig. S11). These species tend to have smaller body sizes and higher LD1 values, indicating characteristics more aligned with highly specialized hypocarnivores. While we did not conduct specific tests on this hypothesis (due to the smaller sample size), we speculate that species with greater specialization towards hypocarnivory might display a similar positive age-dependency dynamic as observed in hypercarnivorous species. Balisi *et al*. (2018) demonstrated that species at both extremes of diet specialization exhibit similar macroevolutionary trends, characterized by shorter durations. This suggests that species at both ends of the specialization spectrum may share similar evolutionary dynamics.

We expected to find a stronger signal of negative age dependency for hypercarnivores, but our results suggest the exact opposite (Fig. 3). Negative age-dependent extinction is typically attributed to population size and geographical range (area) dynamics (see also Saulsbury *et al*., 2023 for a mechanistic model). According to this explanation, newly formed species typically originate within a highly constrained geographical range and possess small population sizes (Vrba, E. S. & DeGusta, 2004; Rosenblum *et al*., 2012). However, as these species age, their geographical ranges and population sizes tend to expand (Miller, 1997; Liow & Stenseth, 2007; Kiessling & Aberhan, 2007), resulting in a lower vulnerability to extinction (Payne & Finnegan, 2007; Foote *et al*., 2008; Jablonski, 2008). Although hypercarnivores typically have lower population densities (Van Valkenburgh *et al*., 2004), which presumably lead to a higher extinction rate (Purvis *et al*., 2000), in particular when new species emerge, they are also expected to have large geographical distributions that typically decrease the probability of extinction (Van Valkenburgh *et al*., 2004; Payne & Finnegan, 2007; Foote *et al*., 2008; Jablonski, 2008, but see Balisi *et al*., 2018). A larger geographical distribution and large body size might indicate a higher dispersal rate (Hoekstra & Fagan, 1998) which would allow them to quickly escape the initial stages of a higher probability of extinction and induce an extinction regime dictated by other factors that more strongly affect older species.

Even though positive ADE may be an artifact produced by an increase in the chance of pseudo-extinction as species get older (Doran *et al*., 2006), positive age-dependent extinction patterns have been attributed to general evolutionary ratchet mechanism leading to “species senescence” and lower evolvability (akin to the ratchet process described by Muller, 1964), and the evolution of specialization (Pearson, 1995). Those hypotheses typically evoke evolutionary changes happening within a given species, not among species, as the most important underlying factor. To explain positive age-dependent extinction, some authors have emphasized specific scenarios such as the absence of evolutionary novelty (Condamine *et al*., 2021), and the inability to adapt in response to a changing environment (Jouault *et al*., 2022).

Hypercarnivore adaptations which include the simplification or loss of dental structures (Holliday & Steppan, 2004; Van Valkenburgh, 2007) exemplify Dollo’s law. According to this law, once a structure is lost (or changed to another state), it is unlikely to be regained (or reverted to the ancestral state) (Holliday & Steppan, 2004). Using a phylogenetic approach, Slater (2015) showed that for Canidae, there is not a single reversal from hypercarnivores to non-hypercarnivores. While the evolution of larger body size and carnivorous adaptations may confer advantages at the individual level (Griffiths, 1980; Carbone *et al*., 2007; Rasmussen *et al*., 2008), it can lead to a macroevolutionary ratchet effect (Van Valkenburgh *et al*., 2004, but see Balisi & Van Valkenburgh 2020). This effect causes hypercarnivory to evolve repeatedly and independently among unrelated lineages (Van Valkenburgh, 2007) with reversals being rare or non-existent (Van Valkenburgh *et al*., 2004; Holliday & Steppan, 2004; Slater 2015). Although the macroevolutionary ratchet mechanism might be seen as a higher-level selection mechanism, the evolution of hypercarnivory (here described as a categorical variable) might also lead to a scenario where species become progressively (through anagenesis) more “hypercarnivore” as they age, which would lead to a positive age-dependent extinction pattern. Holliday & Steppan (2004) found that hypercarnivores faced more pronounced limitations in their subsequent morphological evolution compared to less specialized forms. Their research strongly indicates that as taxa evolve towards hypercarnivory, the degree of morphological flexibility undergoes a significant reduction relative to a more generalist form. Consequently, specialists like canids hypercarnivores may exhibit reduced evolvability (Kirschner & Gerhart, 1998; Wagner, 2008), narrowing their capacity to respond to environmental changes and selection pressures over evolutionary timescales.

It is worth noting that, as pointed out by Finnegan *et al*. (2008), the changes in extinction rates as lineages age could result from an overall change in the “fitness” within each lineage, either through anagenesis or via higher-level selection (in a broad sense) against “less fit” lineages. As pointed out above, most of the discussion on the effect of hypercarnivory in extinction probabilities centers around mechanisms of higher-level selection (Van Valkenburgh *et al*., 2004, but see Balisi & Van Valkenburgh 2020). Much less is known about how the evolution of such traits within a given species might impact its probability of extinction as the species ages. Contrasting the lower and higher-level mechanisms through the lens of different species pools (phylogenetic vs ecological) might help our understanding of the likelihood of ratchet and specialization mechanisms being the underlying mechanisms producing positive ADE for hypercarnivores. Unfortunately, the fossil record is not complete enough to allow us to track such morphological trajectory within species for several species, not allowing such anagenetic mechanism to be properly studied.

That said, the temporal trend of increase in body size and specialization described for the Canidae clade (Van Valkenburgh *et al*., 2004; Silvestro *et al*., 2015), the suggestion that hypercarnivory leads to a higher extinction rate (Van Valkenburgh *et al.,* 2004; but see Balisi & Van Valkenburgh 2020), and the lack of reversion to a non-hypercarnivores state (Slater, 2015), seem to undermine the anagenetic mechanisms as the only one potentially relevant explanations to the positive ADE seen in hypercarnivores described here. As the diversification unfolds and new hypercarnivore species emerge, especially if they exhibit an even higher degree of hypercarnivory compared to their ancestral species (Van Valkenburgh *et al*., 2004), the effect of hypercarnivory on extinction risk would if anything, increase its risk of extinction for newly produced species. Because those would also be the younger species in the pool of species, this macroevolutionary effect of hypercarnivory would tend to produce either a negative ADE or AIE. This suggests that other mechanisms or a combination of these mechanisms are at play here. Pearson (1995) reaches similar conclusions about the implausibility of a “senescence” mechanism as a potential explanation for positive ADE in planktonic foraminifera, trilobites, condonts, and graptolites, raising the possibility that in fact factors extrinsic to the species themselves, such as environmental change, might strongly contribute to generating positive ADE patterns.

We suspect that ecologically driven mechanisms combined with the macroevolutionary ratchet might also be at play for Canidae species in North America. Recent unpublished preliminary analysis by Quental *et al*., suggests that as hypercarnivore species age, they become progressively more “crowded” in the eco-morpho-space. This increase in crowding could be interpreted as an increase in competition pressure as species age, which could represent an ecological mechanism for an increase in extinct risk as species become older and together with a ratchet mechanism could explain the positive ADE. To some extent, this is similar to the “evolutionary stasis hypothesis” discussed by Cid *et al*. (2024) which states that under a scenario of morpho-ecological stasis (reviewed and discussed by Eldredge *et al*., 2005), species’ long-term stability (e.g., the pattern expected under a model of punctuated equilibrium) might lead to a competitive disadvantage of older species when compared to younger species. This idea also resonates with the work of Pearson (1995) who suggests that factors extrinsic to the species itself might play a very important role in determining positive ADE. Hence, we advocate that an ecological perspective should be explored to, in addition, to infer ADE patterns as done here, further investigate the underlying mechanisms of age-dependent extinction in other lineages.

### Methodological implications

In windows characterized by shifts in the extinction rate, we observed a range of different patterns in age-dependent extinction, from very similar signals to their constituent windows to quite a different one. The window Borophaginae 01+02 exhibits a pattern similar to other windows from this subfamily (Fig. 3). The window Non-hypercarnivores 03 results in a clear negative age-dependent dynamic, while one of their constituent windows (the one long enough to be analyzed individually) show a signal more akin to AIE (Fig. 3). It is worth noting that apart from including a shift in rate, this window is the only one that reaches the present. It is possible that edge effects and/or the shifts themselves interfere with the estimates, even though the method was supposedly designed to deal with the former and be quite robust to violations of rate change (Hagen et al 2015). The window Canidae 01+02, showed a very distinct pattern when compared is two constituent windows. While windows without the shift showed evidence for age-independent extinction or distribution skewed towards more negative values, the Canidae 01+02 window displays a distribution skewed towards more positive values. Hence, although Hagen *et al*. (2018) suggested, based on simulations, that ADE inference might be robust to changes in background extinction rate, we argue that the ADE inference might be more sensitive than previously thought, and affected by an unexpected mechanism, of the rate of speciation.

At the genus level, speciation plays a crucial role in defining the persistence of genera which depends not only on their extinction rate but also on the rate of speciation of their constituent species (Raup, 1978). When analyzing ADE at a given taxonomic level, changes in the rate of origination (be it origination at the genus, or speciation at the species level) were not considered to be relevant (Januario & Quental, 2021), but we suspect that those changes in speciation might in fact be relevant when shifts in extinction are also present. When both speciation and extinction change through time and we use a longer time window that encompasses shifts in both rates we might amplify the effect of combining two pools of species with very different longevities. If speciation remains constant, the pool size is only affected by changes in extinction but if speciation also changes, we are more likely to combine different pools of species with very different characteristics, creating a more heterogeneous pool, resulting in different ADE signals. In fact, the simulations conducted by Hagen *et al*. (2018) suggest that changes in extinction rate do not alter ADE inferences using constant speciation rates. We note that changes in speciation rate *per s*e are not problematic for ADE inferences when extinction remains constant or we restrict the analysis to windows of constant background extinction, and this might be why previous studies did not fully appreciate the potential effect of shifts in speciation. Here the species pool belongs to the same extinction regime and the shift in speciation only changes the “number” of species added to this pool. Future works using simulations to analyze windows with shift extinction under scenarios of constant and changing speciation would be conducted to test the robustness of the ADE model under different violations.

## Acknowledgments

We would like to acknowledge Lee Hsiang Liow and Graham Slater for their valuable comments and insights during the course of this work. We are also thankful to Daniele Silvestro for assistance with PyRate and to Lucas M. V. Porto for providing the fossil occurrence database. Additionally, we extend our appreciation to Daniela Munhoz Rossoni, Mathias Mistretta Pires, and Renan Maestri for their suggestions and for being part of the committee for SGN’s thesis defense. The contributions of paleontologists who have studied American mammals, as well as the contributors to the Paleobiology Database and NOW database, are invaluable. This work was funded by FAPESP (#2021/04258-9) to SGN and FAPESP (#2021/06780-4) to TBQ. This is Paleobiology Database publication number XXXXX.

